# Attentional Focus Modulates Automatic Finger-tapping Movements

**DOI:** 10.1101/2020.03.26.009142

**Authors:** Xilei. Zhang, Xingxun. Jiang, Xiangyong. Yuan, Wenming. Zheng

## Abstract

The majority of human behaviors are composed of automatic movements (e.g., walking or finger-tapping) which are learned during nurturing and can be performed simultaneously without interfering with other tasks. One critical and yet to be examined assumption is that the attention system has the innate capacity to modulate automatic movements. The present study tests this assumption. Setting no deliberate goals for movement, we required sixteen participants to perform personalized and well-practiced finger-tapping movements in three experiments while focusing their attention on either different component fingers or away from movements. Using cutting-edge pose estimation techniques to quantify tapping trajectory, we showed that attention to movement can disrupt movement automaticity, as indicated by decreased inter-finger and inter-trial temporal coherence; facilitate the attended and inhibit the unattended movements in terms of tapping amplitude; and re-organize the action sequence into distinctive patterns according to the focus of attention. These findings demonstrate compelling evidence that attention can modulate automatic movements and provide an empirical foundation for theories based on such modulation in controlling human behavior.

## Introduction

The majority of human behaviors are composed of automatic actions ^1–3^. Other than reflexive actions (e.g., the eye-blink reflex evoked by a startling sound), which are natural and can severely interrupt the ongoing task, automatic movements in this study are operationally defined as movements learned during nurturing and can simultaneously be performed without interfering with other tasks ^4–7^. Requiring little attentional control, these automatic actions offer the advantage of allowing us to devote our limited attentional resources to other important matters. For example, we can drink a cup of coffee while immersed in deep thinking without worrying about the muscle contractions and relaxations used to execute these movements. However, this does not mean that automatic movements are disconnected from our attention. On the contrary, attention has long been conceptualized as a system of great significance that modulates automatic movements ^5,8,9^.

On the one hand, it has been proposed that focusing attention on body movements can disrupt movement automaticity and thus deteriorate the performance of well-practiced skills (i.e., constrained action hypothesis ^10,11^). Taking a vertical jump-and-reach task as an example, focusing attention at the fingertips versus the rungs to be touched leads to decreased maximum vertical jump height ^12^. However, it should be noted that these studies involved a hidden common confounding factor. Although these actions are generally deemed skillful and automatic, most are guided by deliberate goals. For example, participants were required to jump as high as possible ^12^, swim as fast as possible ^13^, or throw a dart as accurately as possible ^14^. With specific goals to fulfill, it is natural to assume that participants would deliberately control certain aspects of their movements (e.g., force, timing, speed, coordination). As a result, such studies are incapable of elucidating the complex relationship between attention and automatic movement, given it is impossible to conclude that such actions are not entirely voluntary.

On the other hand, attention has long been conceptualized as the mediator in translating deliberate goals (e.g., drinking coffee) into specific body (muscle) movements (i.e., Normal-Shallice theory ^5^). As one of the core proposals, attention is given an innate potential to modulate automatic movements by facilitating attended actions while inhibiting unattended actions ^5^. To our surprise, this proposal has never been empirically tested (see Discussion). Recently, one neuroimaging study ^15^ found that evidence indicating the interaction between attention and automatic movements is absent for healthy participants (but present for a group of Parkinson’s disease patients). For this study, the authors trained a visually cued keypress movement to automaticity at first (see ^16,17^ for debates on this paradigm in training movement automaticity) and examined neural dynamic changes when attention was directed back to movement. In the healthy control group (as compared to Parkinson’s disease patients), however, data revealed that attention modulates activity and connectivity in those regions responsible for goal-directed, controlled movements (the dorsolateral prefrontal cortex, anterior cingulate cortex, and rostral supplementary motor area) ^15,18^, but not in the region responsible for well-learned, automatic movements (the striatum) ^7^. Though worthy of further scrutinization using more sensitive analyses (e.g., pattern analysis), the non-significant effect of attention on neural activities in the striatum poses a challenge for determining whether attention truly modulates automatic movements of healthy people free of movement disorders.

The present study re-investigates the interaction between attention and automatic movements. Crucially, we did not ask participants to perform automatic movements under any specific goals, given the aforementioned issue of previous studies. Moreover, the automatic movement chosen for this study was not one newly learned in the lab. During both the entrance exam and the main task (see Methods section), participants were required to repetitively and sequentially tap their fingers at a usual, comfortable pace. Even when attention was directed toward these movements, the lack of a specific goal denied the need for participants to try and control their tapping movements. To quantify the trajectories of tapping fingers, we used a new marker-less video-based pose estimation technique ^19^. In the present study, we manipulated attention by asking participants to focus their attention on the movements of either all fingers or a single finger (the index or middle finger), and contrasted these movement-focused conditions with a reference condition in which attention was directed away from the tapping fingers. Given everyday movements are hard to quantify in the laboratory due to techniques used ^20^, evaluation of the direct relationship between attention and automatic movements is a blank to be filled. In the present study, we aimed to fill this blank for the first time, based on methodological improvements introduced above.

To fully elucidate how attention modulates automatic finger-tapping movements, we examined the following three predictions. First, we tested the core proposition of the constrained action hypothesis ^10,11^ that attention disrupts movements’ automaticity. For a sequence rhythmically tapped in perfect automaticity, we assumed that trajectory of finger movements, as well as phase lag between fingers, would be duplicated cycle by cycle, leading to ceiling inter-finger/inter-trial coherence. However, when attention is directed to movement, we predicted a decrease in temporal coherence relative to the reference condition when attention was directed away from movements. Secondly, using tapping amplitude as a measure, we tested a core proposition from the Norman-Shallice theory. This theory, focusing on how attention translates goals into behavior, proposes that attention modulates automatic movements by facilitating attended actions while inhibiting unattended actions. Our primary interest was whether the tapping amplitude would increase for attended fingers while decreasing for unattended fingers. Finally, we performed a pattern analysis on the tapping trajectories of all fingers to examine the prediction that attention focused at different fingers would create distinct patterns of finger-tapping movement. The above predictions were tested in three experiments. Experiments 1 and 2 directed participants’ attention to the respective fingers using two different paradigms, and we expected similar findings in both. For the control, experiment 3 directed attention toward the conceptual label, rather than movement, of respective fingers, and we expected no modulating effect of attention.

## Methods

### Participants

Sixteen college students (7 females) with a mean age of 24.44 ± 0.48 years [hereafter the value following the symbol ‘±’ represents one standard error of the mean (SEM)] took part in experiments 1, 2, and 3 successively over three days spanning eight weeks. The sample size was determined according to previous publications ^21–23^. Procedures and protocols for this study adhered to tenets of the Declaration of Helsinki and were approved by the review board of the School of Biomedical Sciences and Engineering, Southeast University, China. All participants were naive to the purpose of the experiments, gave prior informed written consent, and were debriefed after each experiment.

### Materials

In each experiment, participants were asked to tap their right-hand fingers on a pad fixed to the table and to rest their thumbs at an anchor position marked by a small rubber patch on the pad. Participants’ finger-tapping behaviors were recorded by a full HD-camera (TC-UV8000, 20X-zoom, resolution = 1920×1080, sampling rate = 60 Hz) facing the right hand and placed on the table at a distance of 30 cm from the anchor. When participants were informed to tap their fingers using a pure tone, a visual cue was simultaneously presented on the screen in an isolated area. This area was shielded from the sight of participants but continuously monitored by another HD-camera (TC-980S, 12X-zoom, 1920×1080, 60 Hz). We used a video broadcasting station (model: iRBS-V8, with two video engines) to synchronize videos from these two cameras so we could determine from which frame participants were informed to tap their fingers.

### Procedure

Stimuli were presented using Psychophysics Toolbox extension ^24^ with MATLAB (The MathWorks, Natick, MA). Upon arrival at the laboratory, participants received an entrance exam in which they had to concurrently perform two tasks. In one task of the exam, we required them to tap the fingers (excepting thumbs) of their right hand at their own tempo, ensuring familiarity and comfort with the movements as much as possible. In a simultaneous secondary task, we asked participants to read a text paragraph clearly and loudly. Following prior literature ^6^, we surmised that if one could tap his/her fingers fluently without being interrupted by the secondary task, he/she was regarded as qualified for the subsequent experiment. In fact, participants had no difficulty in passing the exam after some practice.

In experiment 1 (Fig. 1a), we examined whether and how these automatic movements are influenced by the focus of directed attention. In a dual-task paradigm used to check for movement automaticity ^25,26^, participants were required to perform the finger-tapping task simultaneously with a secondary, more exhausting task (i.e., a letter-counting task). This paradigm allows us to evaluate the validity of our manipulation of attentional focus ^27,28^ by measuring the accuracy of the secondary task. Specifically, if participants focused their attention on finger-tapping, the accuracy of the letter-counting task would decrease. The four conditions in experiment 1 included three movement-focused conditions and a reference condition. In the movement-focused condition, attention was directed at the movement of the tapping sequence (sequence-focused condition), the index finger only (index-focused condition), or the middle finger only (middle-focused condition). Under these conditions, participants had to focus their best on movement of the required finger(s), while paying less attention to the letter-counting task. In the reference condition, attention was focused on the sequence of letters (letter-focused condition), and participants were required to count for the appearance of a pre-defined target letter as accurately as possible while neglecting their finger-tapping movements.

**Figure 1.**
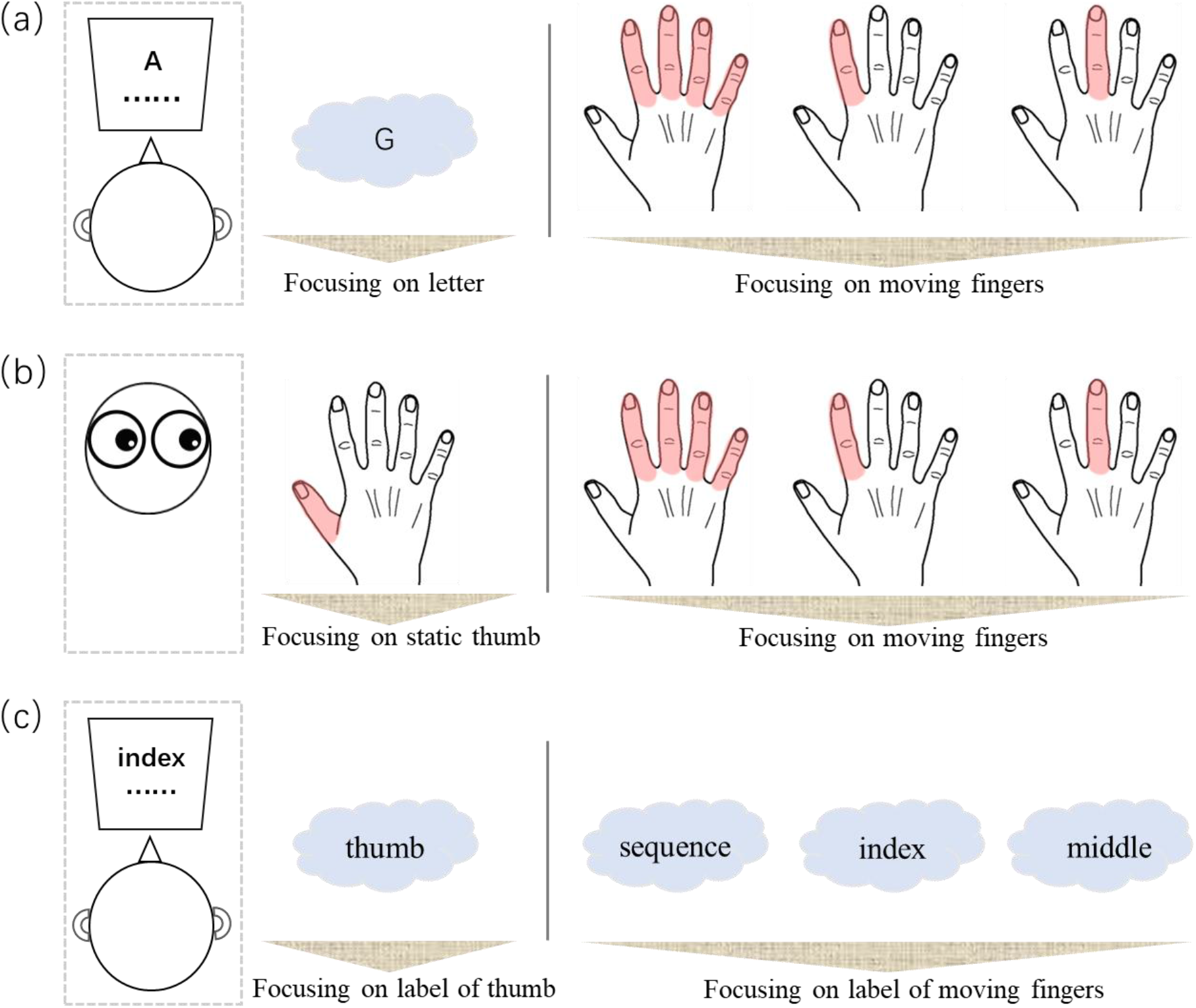
Schematic illustration of the experimental paradigm of (a) experiment 1, (b) experiment 2, and (c) experiment 3. In experiments (a) 1 and (c) 3, a secondary letter-counting or label-counting task was performed simultaneously with the finger-tapping task. The focus of attention was manipulated in each experiment so that either different tapping fingers (indicated by the pink shade on hand) were attended (experiments (a) 1 and (b) 2), or different labels of particular fingers were attended (experiment (c) 3).

At the beginning of each trial, participants were informed of the experimental conditions and target letter (O, G, L, or A, determined randomly). Once their attention was given the required focus, participants pressed the enter key. An initiation beep (500 Hz, 500 ms) indicated participants were to start tapping their fingers until hearing the termination beep (250 Hz, 500 ms). They were advised in advance that any deliberate control of movement was forbidden. During this period, a random sequence of the letters O, G, L, and A were presented at a frequency of 2.5 Hz (stimulus onset asynchrony: 400 ms). Each trial lasted either 6.71 sec (short-duration trial, in which the target letter was presented 3–6 times) or 11.14 sec (long-duration trial, in which the target letter was presented 5–9 times), in order to prevent participants from being able to predict when the trial would terminate. At the end of each trial, participants reported (by pressing keys using their left hands): (1) how many times they saw the target letter, and (2) the amount of attention (ranging from 0 to 100) they had directed to their finger-tapping movements. The next trial began after these responses. A total of 72 trials were segmented into 12 blocks, each corresponding to one condition. In each block, there were three short-duration trials and three long-duration trials. A rest followed every three blocks.

Experiment 2 (Fig. 1b) aimed to replicate the findings of experiment 1 in a single-task paradigm, where participants were instructed to concentrate their efforts on finger-tapping with no secondary task. In line with experiment 1, there were three movement-focused conditions and one reference condition. However, differently and crucially, in the movement-focused conditions, participants were not only told to attend towards, but to forcibly stare at, the particular finger(s), continuously tracking their movements and neglecting the movements of other fingers. In the reference condition, they were told to stare at their static thumbs. The target finger was determined randomly for each trial, with no more than two repetitions. At the end of each trial, participants reported the amount of attention paid to the target finger(s). To save time and alleviate fatigue, the total trial number was reduced to 56, with 7 trials for each duration (short and long) and attention condition. All other aspects were the same as in experiment 1.

Experiment 3 (Fig. 1c) served as a control experiment to examine whether the possible influence of attentional focus on automatic finger-tapping is derived from movement being attended or from its conceptual label embedded in the mind. To this end, we used the dual-task paradigm, as in experiment 1, but replaced the letter-counting task with a label-counting task, in which participants were asked to neglect finger tapping but view a random sequence of finger labels displayed on the screen and count for the target labels. The target label was the “thumb finger” in the reference condition, and the “whole sequence”, “index finger’,” and “middle finger” in three movement-related conditions. To direct attention away from their finger-tapping movements, participants were instructed to look straight ahead at a screen and devote all attention to the counting task. All labels were written in the participants’ native language (i.e., Chinese), and the target label was determined randomly for each trial. At the end of each trial, participants reported: (1) the number of target labels, and (2) the amount of attention they had directed to the movement of related finger(s). Other procedures were the same as those used in experiment 1. There were 56 trials in total, with 7 trials for each duration and attention condition.

### Data Analysis

#### Accuracy of secondary counting task

For experiments 1 and 3, we calculated the accuracy (ACC) of the secondary letter-counting task and label-counting task, respectively, taking ACC as 1 if and only if the reported number of targets equalized with that of the actual number of targets. Otherwise, we took ACC as 0. In the algorithm, missing and false alarms had equal influence on accuracy.

#### Video-based estimation of tapping trajectories

Participants’ finger-tapping movements were extracted using a cutting-edge video-based pose estimation algorithm. Specifically, by using DeepLabCut ^19^ running on a Linux (Ubuntu 16.04 LTS) operating system with a graphics processing unit (GPU, NVIDIA TITAN XP, 12 GB of memory), we trained a deep neural network with 50 layers (i.e., ResNet-50) to automatically recognize the coordinates of seventeen key points of interest (Fig. 2a) for each frame during finger-tapping movements (Fig. 2b). In the first step, we selected 25 representative frames for each participant from videos recorded during experiment 1, manually marked coordinates of key points of interest in these frames, and generated a total of 400 marked frames. We then randomly assigned 380 frames to the training dataset and the remaining 20 frames as the test dataset. By using default parameter settings, we trained a ResNet-50 model through 800,000 iterations. To evaluate its recognition performance, we calculated the mean difference (in pixels) between the manually marked coordinates and predicted coordinates. It was 3.19 ± 0.07 pixels for the training dataset and 4.65 ± 0.90 pixels for the test dataset, a negligible error considering there were 1500 × 1080 = 1,620,000 pixels within each frame. Thus, we applied this model to extract coordinates for each frame, participant, and experiment, and only imported the vertical (y-axis) trajectories of fingers into subsequent analyses, since fingers were tapped mainly in the vertical direction. For preprocessing, vertical trajectories were segmented (from 30 frames before onset of the initiation beep to 90 frames after onset of the termination beep), duration-normalized (the duration of each trial was normalized to the mean duration of short- or long-duration trials using function *interpft*.m), and temporally smoothed (sliding window = 6 frames, equaling 100 ms). Finally, outlier values that deviated the previous frame by a minimum distance of 100 pixels were replaced using the interpolation method (*inpaint_nans.m*). Relative to other approaches quantifying finger movements, for example, those using retro-reflective markers ^29^, force transductors ^30,31^ and cyber glove kinematic sensors ^32^, the marker-less video-based pose estimation technique used in the present study provides an advantage of quantifying movements in high precision while allowing participants to tap their fingers freely.

**Figure 2.**
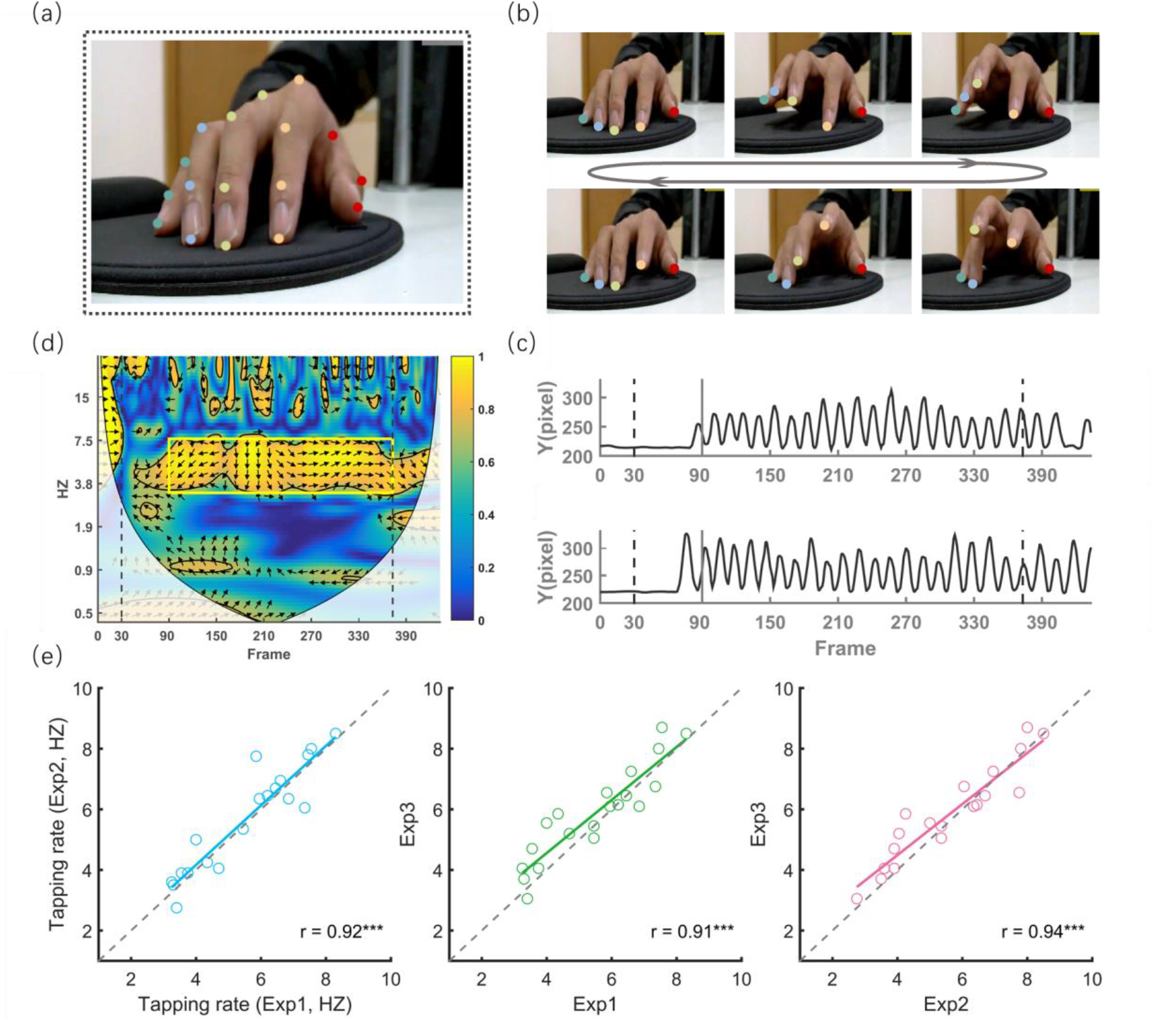
Recording and quantification of automatic finger tapping movements. *(a)* One exemplar frame illustrating the location of 17 key points of interest of the right-hand fingers. For each frame, the coordinates of these points were automatically recognized using DeepLabCut ^19^. *(b)* Concrete frames illustrating changes in the position of key points during repetitive finger tapping movements. For visibility, only key points on the fingertips are shown. *(c)* Y-axis trajectories of the index finger from two exemplar trials. Dotted lines represent the onset of pure tones informing participants to initiate and terminate finger tapping, respectively. The solid line indicates the beginning of the temporal range of interest. *(d)* The 2-D map produced by wavelet transform coherence (WTC) analysis between the above two trials. The color bar indicates the coherence intensity, and the orientations of arrows indicate the circular phase lag at each time-frequency point. The yellow rectangle indicates the temporal-frequency range of interest determined for this representative participant. To exclude the distorting effect of edge artifacts, values in the area masked by translucent white were not included in further analyses. *(e)* Tapping rate is individually different but stable across experiments. Though experiments were performed under different task demands and in separate days, the rate of tapping movements between experiments was highly correlated (Pearson *r* > 0.91).

#### Defining the temporal and frequency ranges of tapping trajectory

The temporal and frequency ranges for our interests were defined as those during which finger-tapping behaviors were automatically performed without deliberate control. First, as the initiation and termination of tapping behaviors inevitably require deliberate control ^5^, we selected the temporal range from 1,000 ms post-onset of the initiation beep to the moment before the onset of the termination beep for further analysis. There were 344 and 609 time points in total for short-duration and long-duration trials, respectively. Second, as automatic finger-tapping behaviors can be deconstructed as repetitive movements of each finger at a constant speed, the frequency range of interest for each participant was defined as the frequency band, which showed the strongest inter-trial temporal coherence. We therefore performed a wavelet transform coherence analysis (WTC, http://www.glaciology.net/wavelet-coherence) for every two trials (Fig. 2c) to produce a time-by-frequency 2-D coherence map (Fig. 2d). As a measure of correlation between two time series ^33^, each coherence intensity out of the coherence map ranges from 0 to 1, with 1 reflecting complete coherence (absolute phase synchrony) and 0 reflecting no coherence (no phase synchrony) at a given time-frequency point. The coherence intensity was then averaged across the selected time range of interest, trial-durations, fingers, and attention conditions. The mean coherence intensity can now be seen as a function of frequency, among which the frequency with the strongest coherence intensity can be located (*findpeaks.m*) and named as the tapping rate. The frequency band corresponding to full width at half maximum (FWHM) centered at the tapping rate was determined as the frequency range of interest.

#### Inter-finger and inter-trial coherence of tapping trajectory

We calculated the inter-finger and inter-trial coherence of tapping trajectory to examine whether and how attention modulates the temporal coupling of finger-tapping movements. For each experiment and condition, mean inter-finger (or inter-trial) coherence was calculated by averaging the coherence intensity within the temporal-frequency range of interest and across trials (or fingertips).

#### Amplitude of tapping trajectory

We calculated the amplitude of tapping trajectories for each fingertip to further test the hypothesis that attention increases tapping amplitude for target fingers and decreases tapping amplitude for non-target fingers. The trajectory of each fingertip was filtered into the frequency range of interest (*filter.m*) and analyzed by Hilbert transform (*hilbert.m*). The absolute value of the resulting signal was averaged within the temporal range of interest, yielding an estimation of the raw tapping amplitude for each condition. For each experiment and fingertip, modulation index I_A_ was computed as I_A_ = (A_attention_/A_reference_)-1, where A_reference_ is the amplitude of the reference condition and A_attention_ is the amplitude of each attention condition. In this way, the I_A_ of the reference condition for each experiment was normalized to 0. Thus, an I_A_ above 0 indicates an excitation effect (i.e., increase in amplitude), while an I_A_ below 0 indicates an inhibition effect (i.e., decrease in amplitude) relative to the reference condition.

#### Pattern analysis of tapping trajectories

Supposing attentional focus indeed interferes with automatic movements, different patterns of finger-tapping behaviors would be observed when the focus of attention is changed. To test this hypothesis, a pattern analysis was performed on the trajectory of all key points from the tapping fingers (a total of 14 key points). First, WTC analysis was performed for each trial between every two key points, resulting in a trial × time (time points in the temporal range of interest) × frequency (40 frequency points) × paired key points 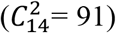 matrix for each participant. The resultant 4-D matrix was then concatenated into a pooled time (trial × time points) × pooled frequency (frequency points × paired key points) 2-D matrix. Second, to reduce dimensional space, we performed a principle component analysis on the pooled frequency dimension, and only retained the top 80 components with the highest variance, which together accounted for over 80% of the total variance. Consequently, the pooled frequency dimension of the 2-D matrix was reduced from 40 × 91 to 80. Third, time points from trials with the same condition were extracted in order and categorized into 6 independent and equivalent datasets. A sliding window method (length = 60 time points, step size = 4 time points, with each step producing a sample containing 60 time points × 80 frequency components) was applied on the pooled time dimension, generating 1188 (or 928) samples for each dataset in experiment 1 (or 2 and 3). Finally, we trained Support Vector Machine (SVM) classifiers (radial basis function kernel, γ = 0.014, as default) for each participant following a six-fold cross-validation approach. In each fold, five datasets were used to train a classifier while the remaining was used for testing. After six folds, all datasets were tested once. To evaluate the performance of the models, we calculated two indices. The decoding accuracy was quantified as the ratio of correctly classified samples to the total number of samples for each condition, while the mean decoding accuracy was calculated as the decoding accuracy averaged across four conditions for each participant and experiment. Beyond the above analyses on temporal coherence and tapping amplitude, analysis on the decoding accuracy provided us the additional advantage to quantitatively evaluate whether and to what extent the two conditions formed the same movement pattern, even if they had comparable mean temporal coherence and/or tapping amplitude.

#### Statistics

Parametric tests used included the one-way and two-way repeated measures analysis of variance (ANOVA), paired-sample t-test, one-sample t-test, and Pearson correlation analysis. Unless specifically denoted, all statistical tests were conducted in a two-tailed manner with a threshold of p = 0.05. All data analyses and statistics were performed using standard and custom scripts adapted to MATLAB and Python.

## Results

### Constant and Fast Tapping Rates between Experiments

As acknowledged, automatic movement operationally refers to those movements that can be performed without interfering with other tasks ^4,5^. From this point of view, if finger-tapping movements were automatic, they would not interfere with the secondary task and would remain stable across experiments, even if conducted on separate days or under different task requirements. Given that each participant’s finger-tapping behavior consisted of a sequence of repetitive movements with which he/she was familiar, the tapping rate of each participant was hypothesized to be stable across experiments. As expected, the tapping rates from three experiments were highly correlated (0.91 < rs < 0.94, ps < 0.001, Fig. 2e). Moreover, participants tapped their fingers at a speed as fast as 5.66 HZ ± 0.35 HZ (ranging from 3.07 HZ to 8.43 HZ). It seems impossible for participants to make deliberate plans and intentionally control their movements at such a fast rate ^34^. No participants reported attempting to control their finger movements when tapping. This converging evidence supports that participants’ finger-tapping movements were automatic and under no intentional control.

### Results of Experiment 1

In experiment 1, a one-way repeated measures ANOVA found a significant effect of attentional focus on both letter-counting accuracy (F_3,45_ = 35.30, p < 0.001, η_*p*_^2^ = 0.702, Fig. 3a) and self-rated amount of attention to movement (F_3,45_ = 204.11, p < 0.001, η_*p*_^2^ = 0.932, Fig. 3b). Relative to the reference condition, post hoc tests revealed that directing attention to movements led to an increase in self-rated amount of attention to movement (all t_15_ > 14.95, ps < 0.001, Cohen’ s *d* > 6.33) and decrease in letter-counting accuracy (all t_15_ < −6.06, ps < 0.001, Cohen’ s *d* > 1.71). Crucially, these two measures correlated significantly for all three movement-focused conditions (−0.73 < rs < −0.61, 0.001 < ps < 0.006, one-tailed, Fig. 3c). These results provide compelling evidence that attention had been effectively directed toward movement in experiment 1.

**Figure 3.**
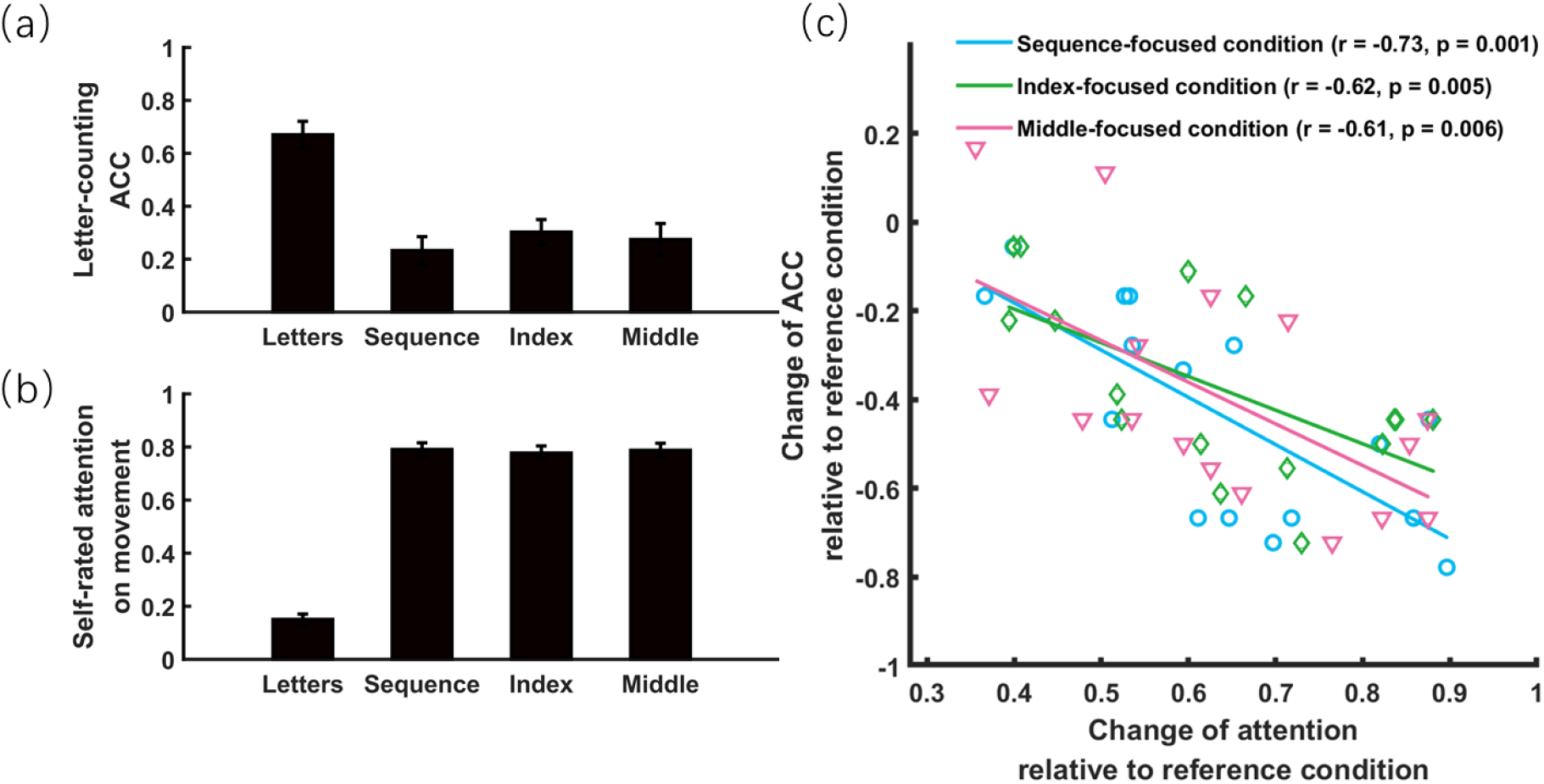
Objective and subjective measures of attention directed to finger-tapping movement in experiment 1. *(a)* Objective measure of attention: accuracy (ACC) of the secondary letter-counting task as a function of attentional focus. *(b)* Subjective measure of attention: self-rated amount of attention directed to movement as a function of attentional focus. *(c)* More attention directed to movement relative to the reference condition (letter-focused condition) led to less attention available for the secondary task, thus possibly contributing to more decrease in ACC of the secondary task. The error bar indicates one SEM.

One of the primary goals of experiment 1 was to examine the prediction derived from the constrained action hypothesis that attention to movement may lead to impaired temporal coherence between tapping trajectory across fingers and across trials. We thus performed two one-way repeated measures ANOVAs on inter-finger coherence and inter-trial coherence, respectively. As shown in Fig. 4a, results revealed a significant effect of attentional focus on both types of coherence measures (F_3,45_ = 7.56, p < 0.001, η_*p*_^2^ > 0.335). Post hoc tests revealed that temporal coherence was strongest in the reference condition (i.e., letter-focused condition; inter-finger coherence: all t_15_ > 1.96, ps < 0.069; inter-trial coherence: all t_15_ > 2.21, ps < 0.043) and weakest in the middle-focused condition (inter-finger coherence: all t_15_ < −3.14, ps < 0.007; inter-trial coherence: all t_15_ < −2.08, ps < 0.055). There were no significant differences between sequence- and index-focused conditions (inter-finger coherence: t_15_ = 0.45, p = 0.661; inter-trial coherence: t_15_ = 0.77, p = 0.453). These results clearly indicate that focusing attention at movement reduces the temporal coherence of automatic finger-tapping movements.

**Figure 4.**
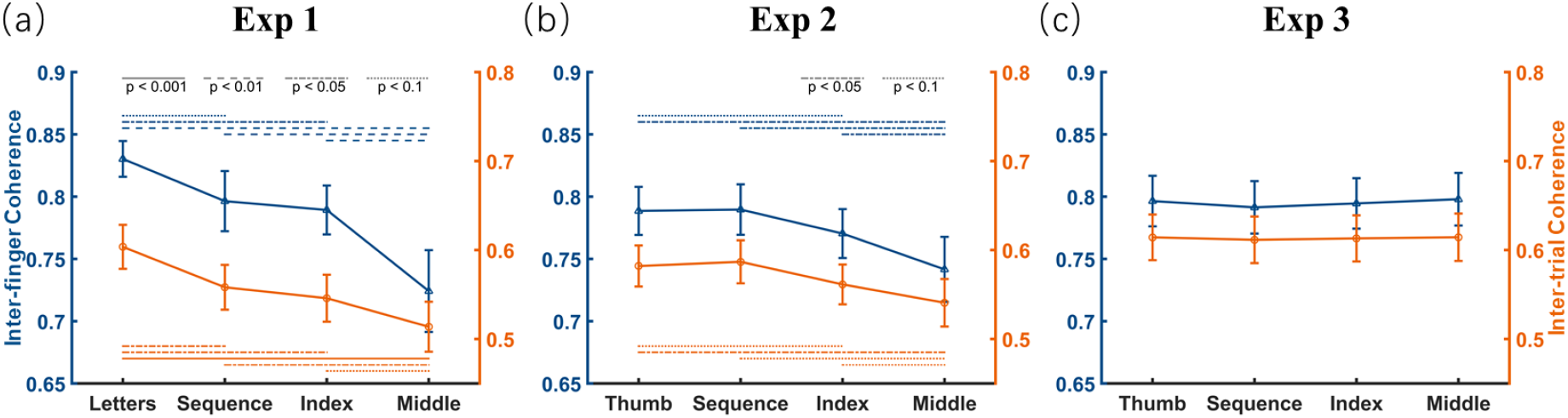
Temporal coherence of finger trajectories varying as a function of attentional focus. Each panel indicates results with respect to inter-finger (shown in blue) and inter-trial (shown in orange) temporal coherence, with x-axis labels indicating four attention conditions with different attentional foci. The error bar indicates one SEM.

Another primary goal of experiment 1 was to examine the prediction derived from the Norman-Shallice theory that attention would facilitate (inhibit) movement of attended (unattended) actions. Following prior literature characterizing the function of the motor system according to the amplitude of tapping movements ^35^, we tested this prediction taking tapping amplitude as the dependent variable. We first examined for each tapping finger to determine whether the I_A_ values in movement-focused conditions were significantly larger than zero using a one-sample t-test, with zero indicating the normalized I_A_ value of the reference condition (i.e., letter-focused condition, see Methods section). For target fingers, expectedly, we revealed I_A_ values significantly above 0 (increased tapping amplitude) for all target fingers in the three movement-focused conditions (all t_15_ > 2.34, ps < 0.033, Fig. 5a). For non-target fingers, however, only the I_A_ value of the little finger was found to have a < 0 tendency (attenuated tapping amplitude, in the index-focused condition: t_15_ = −1.89, p = 0.078, in the middle-focused condition: t_15_ = −2.28, p = 0.038). The I_A_ values of all other non-target fingers in the index-focused and middle-focused conditions did not significantly differ from 0 (all |t_15_| < 1.44, ps > 0.171). These results suggest that attention can facilitate the movement of target fingers and possibly inhibit movement of (some) non-target fingers.

**Figure 5.**
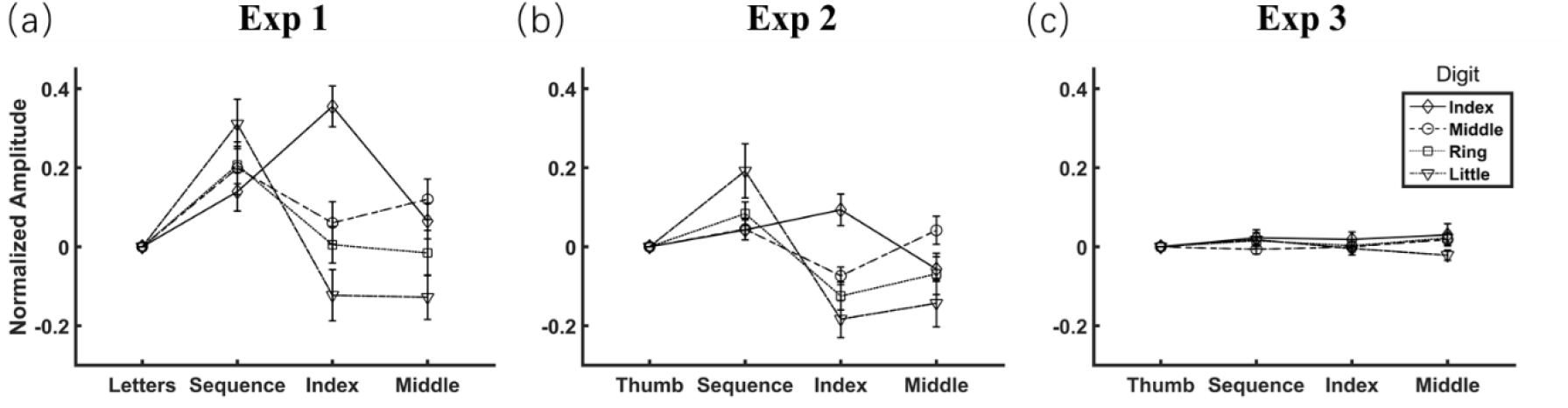
Tapping amplitude of finger trajectories varying as a function of attentional focus. Each panel indicates results with respect to the amplitude for each tapping finger normalized to the reference condition. X-axis labels indicate four attentional focus conditions, with the first label indicating the reference condition whose amplitude had been normalized to 0. The error bar indicates one SEM.

Next, we examined whether the I_A_ values in movement-focused conditions were modulated by tapping fingers. A two-way repeated measures ANOVA on the I_A_ values revealed that the main effects of condition (F_2,30_ = 8.95, p < 0.001, η_*p*_^2^ = 0.374) and tapping finger (F_3,45_ = 8.87, p < 0.001, η_*p*_^2^ = 0.372) as well as the interaction (F_6,90_ = 24.10, p < 0.001, η_*p*_^2^ = 0.616) were significant. To interpret this significant interaction, two follow-up statistical analyses were performed. Firstly, we performed a one-way repeated measures ANOVA on I_A_ values across the three movement-focused conditions for each tapping finger and found significant differences (all F_2,30_ > 6.08, ps < 0.006, η_*p*_^2^ > 0.289). Post hoc tests found a larger tapping amplitude when the index finger was attended than unattended (index-focused condition vs. middle-focused condition: t_15_ = 4.33, p < 0.001; although not significant for sequence-focused condition vs. middle-focused condition: t_15_ = 1.41, p = 0.180). This attention-induced facilitation effect was replicated for the middle finger (sequence-focused and middle-focused conditions vs. index-focused condition: both t_15_ > 2.33, ps < 0.034), and the ring and little fingers (sequence-focused condition vs. index-focused and middle-focused conditions: all t_15_ > 3.26, ps < 0.005). These comparisons further support the effect of attentional focus on tapping amplitude. Secondly, we performed one-way repeated measures ANOVA on the I_A_ values for each movement-focused condition to assess differences in tapping amplitude between fingers, revealing significant results (all F_3,45_ > 4.43, ps < 0.008, η_*p*_^2^ > 0.228). Post-hoc tests showed the tapping amplitude of the little finger increased the greatest among target fingers when attended in the sequence-focused condition (all t_15_ > 2.81, ps < 0.013) and decreased the greatest among non-target fingers when unattended in the index-focused and middle-focused conditions (all t_15_ < −3.25, ps < 0.005). These results suggest that tapping amplitude of the little finger is most sensitive to the effect of attentional focus.

Finally, we performed a pattern analysis to examine whether attentional focus determined the pattern of tapping movements, or, in other words, whether correctly recognizing a particular mode of movement through pattern decoding significantly out-performed guessing (25%). As expected, a one-sample t-test revealed that both the decoding accuracy for each condition (i.e., correct classifications displayed in the diagonal direction of Fig. 6a, all t_15_ > 10.49, ps < 0.001) and the mean decoding accuracy across conditions (64.05% ± 1.87%, t_15_ = 20.91, p < 0.001, Cohen’ s *d* = 5.23, 95% confidence interval, or CI = [3.31, 17.14]) were significantly above chance level. Furthermore, the percentage values in non-diagonal cells (i.e., incorrect classifications) were all significantly below chance level. These results indicate that directing attention to different fingers creates distinct patterns of finger-tapping movements.

**Figure 6.**
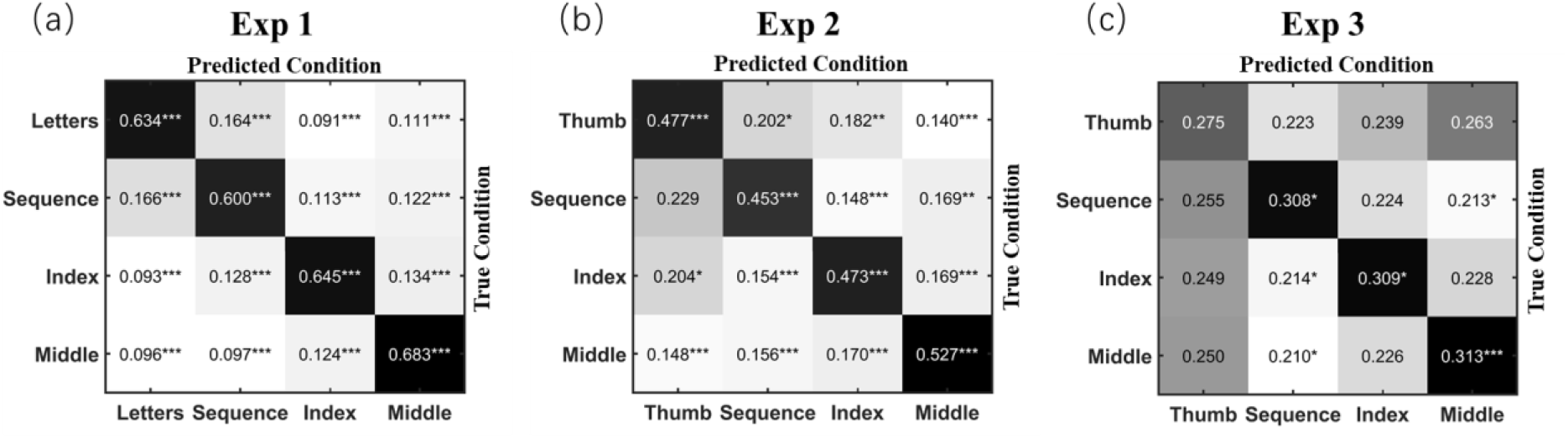
Distinct patterns of automatic movements varying as a function of attentional focus. For each panel and cell, the value corresponds to the ratio of samples of the true condition, which were classified as the predicted condition. Statistics were conducted against the chance level (0.25) by using a one-sample t-test. Darker cells indicate values above chance while whiter cells indicate values below chance. *, p < 0.05, ***, p < 0.001.

### Results of Experiment 2

In experiment 2, we examined whether the effects found in experiment 1 could be observed when participants’ attention was directed toward the target fingers by looking directly at them with no exhausting secondary task. Participants reported that most of their attention (88.27% ± 2.16%) was directed to the target fingers, although they also stated their attention was sometimes distracted by non-target fingers that moved simultaneously with the target finger.

Following the same analyses as experiment 1, the results of experiment 2 first replicated the effect that focusing attention at one’s movement can desynchronize the temporal coherence of automatic finger-tapping movements (Fig. 4b). Specifically, a one-way repeated measures ANOVA found a significant effect of attentional focus on both inter-finger coherence (F_3,45_ = 5.31, p < 0.003, η_*p*_^2^ = 0.261) and inter-trial coherence (F_3,45_ = 3.72, p < 0.018, η_*p*_^2^ = 0.199). Post hoc tests revealed that both inter-finger and inter-trial temporal coherence of the new reference condition (i.e., thumb-focused condition) were larger than those of the index-focused (t_15_ > 1.92, ps < 0.075) and middle-focused conditions (both t_15_ > 2.20, ps < 0.044), but were comparable with those of the sequence-focused condition (both t_15_ < 0.44, ps > 0.668). Consistent with experiment 1, the two indicators of temporal coherence were weakest in the middle-focused condition (inter-finger coherence: all t_15_ < −2.46, ps < 0.027; inter-trial coherence: all t_15_ < −2.05, ps < 0.059), and comparable between the sequence-focused and index-focused conditions (both t_15_ < 1.61, p > 0.129).

With respect to the role of attentional focus on tapping amplitude, we once again compared I_A_ values in the movement-focused conditions with zero, which represented the normalized reference condition (i.e., thumb-focused condition). For the target fingers, as shown in Fig. 5b, one-sample t-tests revealed increased tapping amplitude for the little and ring fingers in the sequence-focused condition (t_15_ > 2.80, ps < 0.014; but for the index and middle fingers: both t_15_ < 1.69, ps > 0.111), and the index finger in the index-focused condition (t_15_ = 2.34, p = 0.034), but not for the middle finger in the middle-focused condition (t_15_ = 1.17, p = 0.260). For non-target fingers, tapping amplitude was attenuated for all non-target fingers in the index-focused condition (t_15_ < −3.30, ps < 0.005) and the index and little fingers in the middle-focused condition (both t_15_ < −1.83, ps < 0.088; but for the ring finger: t_15_ = −1.31, p = 0.209). Collectively, these results suggest that attention can facilitate the movement of (some) target fingers and simultaneously inhibit movement in (some) non-target fingers.

Next, we conducted a two-way repeated measures ANOVA on the I_A_ values, with the condition (three movement-focused conditions) and tapping finger as independent variables, revealing a significant main effect of condition (F_2,30_ = 8.10, p = 0.002, η_*p*_^2^ = 0.351) and a marginally significant main effect of tapping finger (F_3,45_ = 2.27, p = 0.093, η_*p*_^2^ = 0.131); the latter also showed a significant interaction (F_6,90_ = 11.29, p < 0.001, η_*p*_^2^ = 0.429). On one hand, a one-way repeated measures ANOVA indicated significant differences of I_A_ values between the three movement-focused conditions for each finger (all F_2,30_ > 8.03, ps < 0.002, η_*p*_^2^ > 0.349). Post hoc tests replicated the attention-induced facilitation effect (i.e., larger tapping amplitude for the attended fingers versus unattended fingers) for the index finger (sequence-focused and index-focused conditions vs. middle-focused condition: both t_15_ > 3.24, ps < 0.006), the middle finger (sequence-focused and middle-focused conditions vs. index-focused condition: both t_15_ > 3.54, ps < 0.003), and the ring and little fingers (sequence-focused condition vs. index-focused and middle-focused conditions: all t_15_ > 2.14, ps < 0.049). On the other hand, a one-way repeated measures ANOVA showed significant differences between tapping fingers for each condition (all F_3,45_ > 4.54, ps < 0.007, η_*p*_^2^ > 0.232). Consistent with experiment 1, post-hoc tests revealed that amplitude of the little finger increased most when attended in the sequence-focused condition (all t_15_ > 2.81, ps < 0.013) and decreased most when unattended in the index-focused and middle-focused conditions (all t_15_ < −3.25, ps < 0.005). In line with experiment 1, the above results indicate that the effect of attentional focus on tapping amplitude can be observed when participants look directly at the fingers they are instructed to notice.

For pattern analysis, a one-sample t-test yielded an above-chance decoding accuracy for each condition (cells in the diagonal direction: all t_15_ > 5.66, ps < 0.001), and mean decoding accuracy across conditions (48.26 ± 2.61%, t_15_ = 8.91, p < 0.001, Cohen’ s *d* = 2.23, 95% CI = [1.29, 3.14]). In addition, as shown in Fig. 6b, such above-chance decoding performance was observed only for cells in the diagonal direction (i.e., correct predictions), while percentage values in most other cells (i.e., incorrect predictions) were significantly below chance level. These results confirm the findings of experiment 1, indicating that attention directed to different fingers can create distinct patterns of finger-tapping movements.

### Results of Experiment 3

In experiment 3, we directed participants’ attention to the conceptual label instead of the movements of their tapping finger(s). Accuracy of the label-counting task was at a level (72.77% ± 3.82%) comparable with that of the letter-focused reference condition of experiment 1. After the experiment, participants reported that they paid little attention (8.66% ± 1.43%) to finger movements. Thus, participants might direct their main attention toward the conceptual label of target finger(s) and hardly notice their finger-tapping behaviors.

We hypothesized that directing attention towards label and away from movement would have little, if any, impact on automatic finger-tapping movements. To test our hypothesis, we analyzed the data of experiment 3 following the same analyses as experiments 1 and 2. First, a one-way repeated measures ANOVA, taking inter-finger coherence and inter-trial coherence as dependent variables and the focused condition as the independent variable, failed to find a significant effect (both F_3,45_ < 0.69, ps > 0.564, η_*p*_^2^ < 0.045, Fig. 4c). Second, one-sample t-tests showed no significant changes in I_A_ values of the three movement-related conditions relative to zero (|t_15_| < 1.65, ps > 0.120). Third, two-way repeated measures ANOVA on I_A_ revealed that the main effect of condition (F_2,30_ = 0.23, p = 0.795, η_*p*_^2^ = 0.015), main effect of the tapping finger (F_3,45_ = 1.36, p = 0.267, η_*p*_^2^ = 0.083), and their interaction (F_6,90_ = 1.51, p = 0.183, η_*p*_^2^ = 0.092) were all non-significant (Fig. 5c). Finally, we performed a pattern analysis. As shown in Fig. 6c, above-chance decoding accuracy was present only in the three movement-related conditions (t_15_ > 2.45, ps < 0.027), but not in the reference condition (t_15_ = 1.26, p = 0.229). Although a one-sample t-test revealed that mean decoding accuracy was still significantly above-chance (30.12% ± 0.81%, t_15_ = 6.29, p < 0.001, Cohen’ s *d* = 1.57, 95% CI = [0.82, 2.30]), a paired sample t-test revealed a significant decrease relative to the above two experiments (both t_15_ < −6.78, ps < 0.001, Cohen’s *d* > 2.265). As the decoding accuracy for the reference condition did not exceed chance level, it is likely that the partially remaining capability of decoding in movement-related conditions was due to occasionally spreading attention to the movement of respective fingers during the label-counting task, rather than methodological artifacts ^36 37^. Therefore, the above findings confirmed our hypothesis, indicating that it is attention to movement, rather than to conceptual label of respective fingers, that modulates automatic movement.

## Discussion

The present study was, to our best knowledge, the first to demonstrate that attention can modulate automatic movements without deliberate goals. By directing attention to different fingers of an automatic multi-finger repetitive tapping sequence, our main findings were that attention, if focused on movement rather than the conceptual label of different fingers, could lower inter-finger and inter-trial temporal coherence, increase (or decrease) the tapping amplitude of attended (or non-attended) fingers, and create distinct patterns of finger-tapping movements.

A clear distinction should be made between automatic movements we targeted in the present study and reflexive actions (e.g., the eye-blink reflex elicited by a startling sound) that are instinct and prone to severely interrupting the ongoing tasks. Though reflexive actions can also be modulated by attention ^38,39^, they only account for a small proportion of actions in our daily life since they usually emerge in threatening situations. On the contrary, most human behaviors are learned during nurturing and have been well-practiced to automaticity (e.g., walking, talking, or finger-tapping) ^4–7^. As is known, attention is critical for scheduling a variety of our daily movements. Even for movements guided by particular goals, it is the attention, rather than the goals, that directly modulates automatic movements ^5^. From this point of view, we evaluate for the first time in the present study, how attention alone modulates automatic movements when no specific goals are set. Our findings could serve as an indispensable and empirical foundation for theories taking attentional modulation on automatic movements as a prerequisite condition when accounting for the role of attention in complex behaviors such as skill performance and learning ^11,40^ and goal-directed action control ^5^.

First, our findings provide empirical foundation for the core idea of the well-known constrained action hypothesis that attention to movement disrupts movement automaticity ^10,11^. In line with this proposal, we found reduced temporal coherence, indicating disrupted automaticity, in almost all movement-focused conditions compared with the reference condition. The impaired coherence may be accounted for by mismatch on the coordination speed between systems supporting deliberate attention and automatic movements, respectively ^34,41^. As revealed via a task requiring maintaining balance on an unstable surface, attention to foot movement (internal focus condition) leads to slower movement adjustments, while attention directed away from foot (external focus condition) leads to faster movement adjustment ^42^. These results indicate a slower pace when attention is manipulated than in automatic mode. Applying the same logic to the present study, the attention may have actively intervened in the fast-paced automatic movement, leading to timing mismatch (as reflected by decreased temporal coherence) between the attended and unattended fingers.

Our findings also provide an empirical foundation for the renowned Norman-Shallice theory ^5^. This theory focuses on how deliberate goals control automatic movements by directing attention. A critical and yet to be examined assumption is the bi-directional modulation effect of attention upon automatic movements. In other words, attention by itself would facilitate attended actions and inhibit unattended actions. Since daily action is hard to study in the laboratory ^20^, researches testing the Norman-Shallice theory put more effort in addressing the proposed role of the frontal lobe in executive control (inhibitory control and/or conflict monitoring) ^43^, for example using tasks like the Stroop and Wisconsin Card Sort ^44–46^. However, they relatively neglected how attention by itself modulates automatic movements. In the present study, our finding that attention could increase (or decrease) the tapping amplitude of attended (or non-attended) fingers supplies direct and compelling evidence supporting this view. Although a similar effect of attention has been widely proven and accepted in the field of perception ^47^, to our best knowledge, this study is the first to extend this attentional effect to automatic movements.

Results of the pattern analysis provide more insight about the interaction between attention and automatic movements. As revealed, the above-chance decoding accuracy indicates that attention can dramatically reorganize automatic movements into distinctive patterns, even when those movements cannot be distinguished from each other according to their temporal coherence or tapping amplitude. Highlighting the intimate interaction between attention and automatic movements, the pattern analysis findings support both the constrained action hypothesis ^10,11^ and the Norman-Shallice theory ^5^. The dependence of automatic movement patterns varying as a function of attentional focus may be double-edged. On one hand, newly created movement patterns may fail to satisfy previous requirements and consequently lead to deteriorated task performance ^40^. On the other hand, disruption of the previously established movement automaticity is necessary to acquire new motor skills, especially in the early learning stage ^8,9^.

All findings above conclude with the fundamental view that attention can modulate automatic movements, even without specific goals. However, the neural modulations, when attending to automatic movements, have barely been investigated. Prior studies have revealed that the neural dynamics underlying the development of movement automaticity (task expertise) are mainly two-folds. On the one hand, neural activation decreases in a wide variety of motor-related areas (e.g., cerebellum, cingulate motor area, primary motor cortex and supplementary motor area)^15,48,49^ and the attention system (especially the dorsolateral prefrontal cortex, i.e., DLPFC, and anterior cingulate cortex, i.e., ACC ^18,50^). These results suggest that attention becomes less important when there evolves a more efficient neural code for controlling movements. On the other hand, neural activation increases in a critical subcortical region, i.e., the sensorimotor territory of the striatum ^7^, as well as its effective connectivity with these motor-related regions ^15,48^. These results suggest that motor skills are likely encoded across multiple representations within a subcortico-cortical loop ^51^ centered on the striatum.

When attention is directed back to automatic movements, we surmise that the subcortical striatum would be of different neural activity and connectivity with motor-related regions and the attention system. Though prior studies examining this possibility ^15,18^ provided no support for this hypothesis, two critical issues should be considered, pointing out future studies’ direction. At first, the task they used (i.e., initiation of visually cued keypress movements) might have mainly measured dynamics related to deliberate control processes, because initiation of a movement sequence, which usually requires deliberate control ^5^, can hardly be trained to automatic despite the level of training ^16,17^. For this reason, the striatum may not be effectively engaged in related tasks. Alternatively, the paradigm used in the present study may provide a promising solution. On the other hand, the fact that they did not find differences in the striatum does not necessarily mean differences do not exist. For this reason, future studies are encouraged to employ more sensitive analyses (e.g., pattern analysis) to delineate neural dynamic changes, if any, in the striatum and other related regions.

Future studies are also encouraged to compare the neural dynamic changes between the healthy and motor deficit (e.g., Parkinson’s disease) patients, to obtain a better understanding of motor-deficit pathophysiology. Inspired by the Norman-Shallice theory ^5^, continued investigations should also bring goals into consideration ^52^, to further examine whether and how attention translates deliberate goals into automatic movement sequences. Limitations of this study include that we did not track eye movements, which prevented us from evaluating to what extent participants had locked their gaze on the required focus. The fact that we did not counterbalance the order of experiments constitutes another limitation, although the order effects, if any, would decay during the interval (spanning weeks) between experiments.

In sum, we found for the first time that attention can modulate automatic movements without assistance from deliberate goals. Once being directed to perform a sequence of automatic movements, focused attention disrupts movement automaticity, facilitates attended and inhibits unattended movements, and re-organizes the movement sequence into distinct patterns according to the focus of attention.

## Acknowledgements

This research was supported by grants from the National Key Research & Development Program of China (2018YFB1305200) and the National Natural Science Foundation of China (61921004) for W.Z., and Postdoctoral Science Foundation of China (2019M661703) and the Fundamental Research Funds for the Central Universities (2242020R20021) for X.Z.

## Author Contributions Statement

X.Z. developed the study concept and designed the study. Testing and data collection were performed by X.Z. and X.J; X.Z., X.J., and X.Y. performed the data analysis and interpretation under the supervision of W.Z; X.Z. drafted the manuscript, and X.Y. and W.Z. provided critical revisions. All authors approved the final version of the manuscript for submission.

## Additional Information

The authors declare no competing interests.

## Notes

### Competing Interest Statement

The authors have declared no competing interest.

### Summary of Updates

Manuscript revised according to the comments mainly from two anonymous reviewers invited by the journal Scientific Reports.

